# Y-chromosomal microsatellites and expansion from Africa

**DOI:** 10.1101/2023.04.29.538812

**Authors:** Zarus Cenac

**Affiliations:** Department of Psychology, City, University of London, Rhind Building, St John Street, London, United Kingdom, EC1R 0JD

## Abstract

Research has suggested that Africa is the continent from which modern humans dispersed around the world. Some diversities are thought to trace this expansion; they decline as distance from Africa accumulates, and they suggest that the expansion originated within a range of locations in Africa. Previously, a decline was found regarding Y-chromosomal microsatellite heterozygosity. However, this diversity has been noted to suggest a non-African origin and, consequently, not look like it indicates expansion from Africa. Declines have appeared in other variables which are derived from Y-chromosomal microsatellites. These variables include effective population size, expansion time, and time to the most recent common ancestor. The present research inferred if these variables, and some Y-chromosomal diversities, indicate expansion from Africa. Variables which indicate the expansion could help with identifying which area of Africa the expansion started in. This research used variables previously derived from Y-chromosomal microsatellites. These variables were for populations from across the world (effective population size, expansion time, and time to the most recent common ancestor) or only in Africa (haplotype and repeat unit diversities). Regarding populations across the world, San were very high in each variable when considering distance from (southern) Africa. Without San, effective population size and time to the most recent common ancestor did not squarely indicate an African origin, and expansion time placed the origin wholly outside of Africa. When using San, each area was only in Africa. Amongst African populations, whilst both haplotype diversity and repeat unit diversity showed a geographical pattern of declining, only haplotype diversity suggested a completely African area of origin. A southern African origin was broadly indicated when a general estimate of the origin was produced from Y-chromosomal haplotype diversity alongside some other genetic diversities.

## Introduction

Modern humans appear to have gone through a global expansion from the African continent (Ramachandran et al., 2005). Research suggests that they met bottlenecks along the way (Ramachandran et al., 2005).^1^ Diversities vary in the extent to which they represent expansion from Africa (Betti et al., 2012). A decrease of diversity with greatening distance from Africa is consistent with there having been bottlenecks in the expansion (Ramachandran et al., 2005). Decreases are evident, for instance, with autosomal diversity (Ramachandran et al., 2005) and cranial diversity (Manica et al., 2007). However, with some other diversities, no such decline is suggested (Betti et al., 2012). Research with respect to expansion from Africa has concerned Y-chromosomal microsatellites (e.g., Shi et al., 2010). Whilst Y-chromosomal microsatellite heterozygosity does decline (Balloux et al., 2009), it might not denote expansion from Africa (see *Introduction: Worldwide*) (Cenac, 2022). Studies have not merely indicated that Africa is the origin of expansion – they have looked for where the expansion arose from inside of the continent (e.g., Ray et al., 2005; Tishkoff et al., 2009). However, a definitive answer has not been reached with respect to the area of Africa which the expansion set off from (Cenac, 2022). The present study examined whether certain qualities derived from Y-chromosomal microsatellites (de Filippo et al., 2011; Shi et al., 2010) indicate the expansion from Africa. This was examination was done in order to continue assessing where the expansion originated (following on from Cenac, 2022) with the addition of Y-chromosomal microsatellite-derived variables as appropriate.

### Worldwide

A number of variables can be found using Y-chromosomal microsatellites, and these variables are related to distance from Africa (Balloux et al., 2009; Shi et al., 2010). These include heterozygosity (Balloux et al., 2009) and time to the most recent common ancestor (TMRCA) (Shi et al., 2010). Indeed, Balloux et al. (2009) observed that Y-chromosomal microsatellite expected heterozygosity (found from six Y-chromosomal microsatellites) decreases with the increasing of distance from (sub-Saharan) Africa. However, this heterozygosity does not appear to capture the worldwide expansion, because research has found that Y-chromosomal microsatellite heterozygosity i) points to an area of origin for worldwide expansion placed to some extent, or even entirely, in Asia (not Africa), and ii) is predicted better by distance from Asia (specifically India) than by distance from Africa (Cenac, 2022); that research used heterozygosity which Balloux et al. (2009) presented for only six microsatellites (Cenac, 2022). Nonetheless, when using 17 microsatellites, Y-chromosomal heterozygosity would still not seem to be consistent with indicating an African origin (Figure 1).

**Figure 1.**
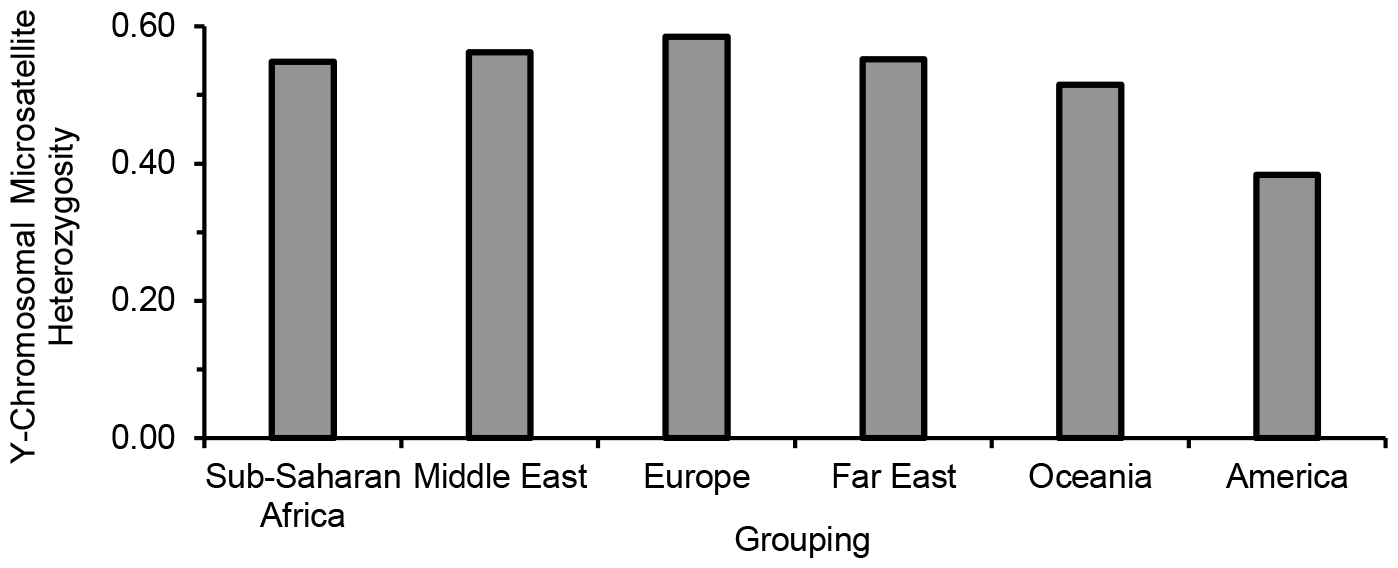
Y-Chromosomal Diversity. *Note*. Y-chromosomal gene diversities (from 17 microsatellites) of populations were found in Xu et al. (2015), and, in the present study, the gene diversities were averaged within regional groupings found in Xu et al. Gene diversity means expected heterozygosity (e.g., Harris & DeGiorgio, 2017). Given comments on p. 143 of Xu et al. (2015), it could very well be that Xu et al. also averaged like the present study did – nevertheless, they do refer to Europe as having the highest gene diversity (and America possessing the lowest gene diversity / heterozygosity) in their study (Europe and America indeed have the respectively numerically greatest and numerically lowest diversity in Figure 1). Regional labels were used from Xu et al. (2015), and are presented on the *y*-axis (in the order given in Xu et al.). A decline in diversity (from relatively higher diversity to lower) fits with expansion – the extent of decline varies with the place employed *as* the origin, with a stronger decline appearing to highlight where the expansion started (Ramachandran et al., 2005; von Cramon-Taubadel & Lycett, 2008). So, Y-chromosomal microsatellite heterozygosity being at its numerical top with Europe (Xu et al., 2015; Figure 1) is not compatible with the expansion from Africa being indicated by this heterozygosity.

Regarding Y-chromosomal microsatellites, levels of TMRCA, effective population size, and expansion time (further into the past) do seem in tune with expansion from Africa (Xu et al., 2015). Indeed, levels are generally greatest in Africa and lowest farther along in the expansion from Africa (Xu et al., 2015). Distance from Africa indeed has correlations (negative, linear ones) with Y-chromosome-derived (through 65 microsatellites) TMRCA, effective population size, and expansion time (Shi et al., 2010). Whilst that study (i.e., Shi et al., 2010) did place median levels of these variables on a map concerning each variable, the study did not distinguish a specific area for the origin of worldwide expansion like has been done for diversities (such as in Manica et al., 2007).

### Within Africa

Examination of whether there are relationships between diversity and distance has happened at the global scale (e.g., Balloux et al., 2009; Betti et al., 2009), or at a smaller scale than globally (e.g., Hunley & Cabana, 2016). For instance, using populations across the Earth, as distance from Africa pushes onward, autosomal microsatellite heterozygosity falls (e.g., Balloux et al., 2009; Prugnolle et al., 2005), and this type of autosomal diversity decreases *within Africa* as distance from *southern* Africa rises (Cenac, 2022). Furthermore, autosomal microsatellite heterozygosity inside Africa may identify where the expansion initiated (Cenac, 2022). As for Y-chromosomal diversity, a local examination could indicate that Y-chromosomal microsatellite heterozygosity *within Africa* is incongruent with expansion from Asia – for populations within Africa, previous research noted a *positive* Pearson correlation coefficient regarding Y-chromosomal microsatellite heterozygosity and distance from Asia, *r* = .70 (if Asia is the origin, then a negative coefficient would be expected), and, furthermore, that research found a *negative* coefficient was possible with distance from within Africa, *r* = -.90 (Cenac, 2022). However, just seven populations were used and statistical significance was not tested for (Cenac, 2022).^2^

### Non-linearity

Preprints have concerned non-linearity (Cenac, 2022, 2023). Unlike autosomal SNP^3^ haplotype heterozygosity, X-chromosomal microsatellite heterozygosity, and cranial shape diversity, both autosomal microsatellite heterozygosity and mitochondrial diversity may be better described as having quadratic relationships with distance from Africa rather than linear declines (although support for a quadratic relationship regarding mitochondrial diversity could arise from climate) (Cenac, 2023). And so, there could be a point in considering non-linearity with respect to distance (e.g., from Africa) and Y-chromosome-derived haplotype or repeat unit diversities, TMRCA, effective population size, and expansion time.

### Origin

There are estimates out there regarding where the expansion emerged from (Ramachandran et al., 2005; Ray et al., 2005). For example, research had sought a general estimation concerning the part of Africa the expansion set out from by using *peak points* – regarding diversity, a peak point is the geographical place from which diversity declines most strongly (numerically) as distance increases (Cenac, 2022). That research arrived at a general estimate for the origin (southern Africa) by seeing where peak points for variables are situated in general, and also by ascertaining a centroid from peak points (Cenac, 2022). However, it might be better to take into account the overall *distribution* of relationships between distance and a variable (e.g., genetic diversity) across potential origins (like in Luca et al., 2011, Ramachandran et al., 2005, or Tishkoff et al., 2009), and then combine these distributions across variables. This was, essentially, carried out in the present study, but with ranks – the ranking of potential origins (see *Method*).

### Objectives

Preceding research has marked out a geographical range of locations (a geographical *area*) in which the *origin* of the expansion could be (e.g., Manica et al., 2007). Origin areas did not appear to have been marked out previously (like in Manica et al., 2007) using TMRCA, effective population size, and expansion time with respect to Y-chromosomal microsatellites or otherwise. Therefore, the present study estimated origin areas using those variables. To do that, this study used TMRCA, effective population size, and expansion time presented in Shi et al. (2010) for populations across the Earth; Shi et al. (2010) found these variables through using 65 Y-chromosomal microsatellites. Therefore, the data used in the current study was from a greater number of Y-chromosomal microsatellites than the six used in Balloux et al. (2009). Following on from previous research concerning diversity (in which an African origin was indicated, but not an additional origin outside of Africa) (Manica et al., 2007), this study explored if there may be more than one origin indicated for Y-chromosome-derived TMRCA, effective population size, and expansion time. As for diversity within Africa, research regarding Y-chromosomal microsatellite diversity has used, and presented, diversity for 26 populations in Africa (de Filippo et al., 2011) (i.e., more than seven populations); the present study utilised repeat unit diversity and haplotype diversity from de Filippo et al. (2011) to test if diversities correlate with distance within Africa,^4 5^ and also to estimate origin areas. Moreover, given prior research (Balloux et al., 2009; Cenac, 2022; Manica et al., 2007), using these diversities, it was inferred if there may be more than one expansion signal – one from within Africa (i.e., perhaps consistent with expansion from Africa, like autosomal microsatellite diversity in Cenac, 2022), and one into Africa. Moreover, it was seen if relationships between geographical distance and Y-chromosome-derived variables (TMRCA, effective population size, expansion time, haplotype diversity, and repeat unit diversity) are represented better as being quadratic than linear. Ultimately, across types of genetic diversities (Balloux et al., 2009; de Filippo et al., 2011; Pemberton et al., 2013), the region of Africa which the expansion spread from was surmised using a ranking-based method referred to above (and described in more detail below).

## Method

### General

Variables were found in previous research, be they regarding the Y chromosome (de Filippo et al., 2011; Shi et al., 2010), the autosome (Balloux et al., 2009; Pemberton et al., 2013), mitochondria, or the X chromosome (Balloux et al., 2009). For analyses, this study utilised R Version 4.0.5 (R Core Team, 2021) and Microsoft Excel. Atypical datapoints were more than 3.29 (in *z*-scores of residuals) away from their predicted value (e.g., Field, 2013) in the present study.

### Y-chromosomal microsatellite data

The current research used variables regarding the Y chromosome (TMRCA, effective population size, and expansion time) displayed in Shi et al. (2010), which Shi et al. (2010) calculated through 65 Y-chromosomal microsatellites. Some population names presented in Shi et al. (2010) were, to various extents, different to ones presented in Balloux et al. (2009), but were similar enough that it was assumed that they refer to the same populations (particularly as both Shi et al. and Balloux et al. used HGDP-CEPH data). So, distances in Cenac (2022) regarding populations in Balloux et al. were applied to the populations used in Shi et al. (e.g., when calculating BICs). This left out one population used by Shi et al. (Bantu of South Africa), which Balloux et al. (2009) did not present Y-chromosomal diversity for as Balloux et al. refer to not including, in their analysis of HGDP-CEPH data, two Bantu populations in the south – the TMRCA, effective population size, and expansion time for that population were omitted in the present study. Shi et al. (2010) combined Colombian and Mayan populations. And so, whilst Shi et al. (2010) used 51 datapoints (in their correlation analysis), the present research used 50 of their datapoints in analyses. Additionally, this study employed two Y-chromosomal microsatellite measures (repeat unit diversity and haplotype diversity) presented in de Filippo et al. (2011) which de Filippo et al. calculated using 11 microsatellites in 26 African populations. With respect to the populations for which haplotype and repeat unit diversities are presented in their study, de Filippo et al. (2011) refer to their own study and also cite Berniell-Lee et al. (2009), Coelho et al. (2009), de Filippo et al. (2010), Gomes et al. (2010), and Tishkoff et al. (2007).

### Origin

The Bayesian information criterion (BIC) was used for estimation of the area of origin, like in preceding research (e.g., Betti et al., 2009; Manica et al., 2007). As before (Cenac, 2022), a variety of locations in previous studies (99 in Betti et al., 2013; 32 in von Cramon-Taubadel & Lycett, 2008) functioned as origins. Following procedures outlined before (Manica et al., 2007), a BIC was calculated for each origin used, and, after the lowest BIC was identified, the area of origin was assembled from that origin in tandem with any origins whose BICs were within four BICs of the BIC which was lowest. BICs were determined concerning geographical distance and i) TMRCA, ii) effective population size, iii) expansion time, iv) repeat unit diversity, and v) haplotype diversity. The calculation of BICs was done in Microsoft Excel through a formula provided in Masson (2011), or in R Version 4.0.5 (R Core Team, 2021). As before with analysis of autosomal microsatellite diversity inside Africa (Cenac, 2022), origins in Africa (99 coordinates in Betti et al., 2013) were used when BICs were determined regarding diversity within Africa in the present study. This applied to Y-chromosomal microsatellite repeat unit and haplotype diversities. Additionally, for the within-Africa analysis, coordinates (a potential waypoint which Tishkoff et al., 2009, estimated for the exit from Africa)^6^ were used in the present study as an origin in BIC calculations. This origin was employed in order to suggest if origin areas are unique to Africa.

When estimating the origin, it can be important to consider whether correlation coefficients are positive or negative – regarding diversity, an origin *should* have a negative correlation coefficient (e.g., Cenac, 2022); this was applied (specification of coefficients being negative) when estimating origin areas. The origin delivering the lowest BIC (amongst origins with negative correlation coefficients) can be called the peak point (Cenac, 2022).

### Two origins?

Previous research has used BICs to examine if there may be two origins (one inside Africa, and one outside) – distance from an additional origin (outside of Africa) was added to the strongest model for predicting diversity (i.e., a model which included distance from the peak point, which was inside Africa), and it was seen if using these distances from the additional origin improved the ability to predict diversity (presumably using the within-four-BIC procedure) (Manica et al., 2007). An improvement would suggest *an* origin outside Africa (Manica et al., 2007). That method was applied, in the present study, regarding TMRCA, effective population size, and expansion time when Africa had the peak point – starting with the model with distances from the peak point, it was seen whether adding distances from any of the origins outside Africa (coordinates from von Cramon-Taubadel & Lycett, 2008) led to a stronger prediction of the Y-chromosome-derived variables (and therefore support for an expansion starting outside of Africa). The method (Manica et al., 2007) was also followed regarding Y-chromosomal diversity – after having found the strongest model for predicting Y-chromosomal diversity (i.e., a model with distances from the peak point), it was seen whether adding distances from the aforementioned Tishkoff et al. (2009) waypoint improved on predictions; an improvement may, therefore, suggest expansion from somewhere other than Africa. The fall of diversity (as distance from some location greatens) is indicative of expansion from that location (e.g., Ramachandran et al., 2005; von Cramon-Taubadel & Lycett, 2008), and Y-chromosome-derived TMRCA, effective population size, and expansion time decline as distance from the apparent origin of expansion (Africa) rises (Shi et al., 2010). And so, an additional origin was supported not purely on the basis of BICs, but also by producing a negative correlation coefficient.

### Quadratic or linear

Like beforehand, BICs were used (including applying the requirement of being within four BICs) (Cenac, 2022, 2023) to assess whether adding a quadratic term to linear (term) models resulted in a better ability of geographical distance to predict genetic-derived variables than the best of the linear models. The variables of interest in the present study were the Y-chromosome-derived TMRCA, effective population size, expansion time, and diversities concerning haplotypes and repeat units.

### Estimating the origin across variables

A decrease of diversity over distances from Africa is compatible with expansion from Africa (e.g., Ramachandran et al., 2005; von Cramon-Taubadel & Lycett, 2008). Moreover, for a diversity variable, a stronger decline indicates better candidacy for being the origin of expansion (Ramachandran et al., 2005). Therefore, *r*s and also *R*^2^s (regarding distances from origins and diversity) can reflect the origin of expansion (Ramachandran et al., 2005; von Cramon-Taubadel & Lycett, 2008); in the present study, an origin was ranked higher according to how *low* the resulting correlation coefficient was (so an origin which has *r* = -1.00 would be ranked above an origin with *r* = -.90) or the size of *R*^2^s (the larger the *R*^2^ the lower the rank). For rankings, the present study calculated and used *r*s and *R*^2^s regarding linear relationships and quadratic relationships respectively (some of which had been calculated before in Cenac, 2022) – see below.

Ranks were found using the following variables: i) autosomal microsatellite heterozygosity which Pemberton et al. (2013) determined from Tishkoff et al. (2009) (the same diversities of 106 populations used in Cenac, 2022) and HGDP-CEPH data (diversities of 50 populations used in Cenac, 2023), ii) autosomal SNP haplotype heterozygosity displayed in Balloux et al. (2009) (which Balloux et al., 2009, got from Li et al., 2008, and Li et al., 2008, calculated the heterozygosity from HGDP-CEPH data), iii) mitochondrial diversity from Balloux et al. (2009), which Balloux et al. sourced from the Human Mitochondrial Genome Database (regarding which they cite Ingman & Gyllensten, 2006) and GenBank, iv) X-chromosomal microsatellite heterozygosity from Balloux et al. (2009) which they calculated from HGDP-CEPH data, and v) (given outcomes in *Results and discussion*) the Y-chromosomal microsatellite haplotype diversity shown in de Filippo et al. (2011).^7^

Relationships between distance from Africa and diversity appear to be better shown through quadratic (than linear) trends for autosomal microsatellite heterozygosity and also mitochondrial diversity, but this is not apparent for autosomal SNP haplotype heterozygosity, X-chromosomal microsatellite heterozygosity (Cenac, 2023) and Y-chromosomal haplotype diversity (present study). Therefore, *r*s were used for rankings regarding autosomal SNP haplotype heterozygosity, X-chromosomal microsatellite heterozygosity, and Y-chromosomal haplotype diversity, whilst *R*^2^s were utilised for rankings concerning autosomal microsatellite heterozygosity and mitochondrial diversity.

These variables have the trait of declining regarding distance from Africa (using populations either worldwide or within Africa) (Balloux et al., 2009; Prugnolle et al., 2005; present study) and they align with Africa individually possessing the area of origin (Cenac, 2022; Manica et al, 2007; present study). To find the ranks, correlation coefficients (*r*s) or *R*^2^s were worked out concerning each variable and distances from the 99 Betti et al. (2013) African origins (through the use of distances from Cenac, 2022, 2023, who used the Betti et al. origins, or distances found in the current study which used those origins too – see *Method: Geographical distance*). Between autosomal microsatellite and autosomal SNP haplotype heterozygosities, ranks were averaged at each origin. Then these averaged ranks were themselves ranked (i.e., an average autosomal rank).^8^ For each origin, an average was taken across the average autosomal ranks, and the remaining ranks (mitochondrial, X-chromosomal, and Y-chromosomal haplotype). These overall averaged ranks were then ranked across origins (i.e., ranked from 1 to 99).

As an indication of where the expansion started, previous research calculated a centroid from peak points for: i) autosomal diversity (as a centroid from three peak points), ii) mitochondrial diversity, iii) X-chromosomal diversity, iv) cranial shape diversity (as an average of two peak points) and v) cranial size dimorphism (Cenac, 2022). The optimal way of combining the peak points is not clear. For example, would it have made more sense to indicate the origin by calculating a centroid amongst genetic diversities (genetic centroid), and an average amongst cranial traits (cranial average), and then averaging between the genetic centroid and cranial average? So, for simplicity in the present study (using the rank-based approach), only genetic diversities were focused on when estimating the origin.

## Figures

Figures 2A-2F, 3A-3C, and Figure 4 were updated/modified from Cenac (2022) to reflect the Y-chromosome-derived data used in the present study or ranked origin locations (Cenac, 2022, employed coordinates/locations in Africa represented in Figure 4 of Betti et al., 2013). This meant that, ultimately, Figure 2F matched Figure 5C in Cenac, 2022).

**Figure 2.**
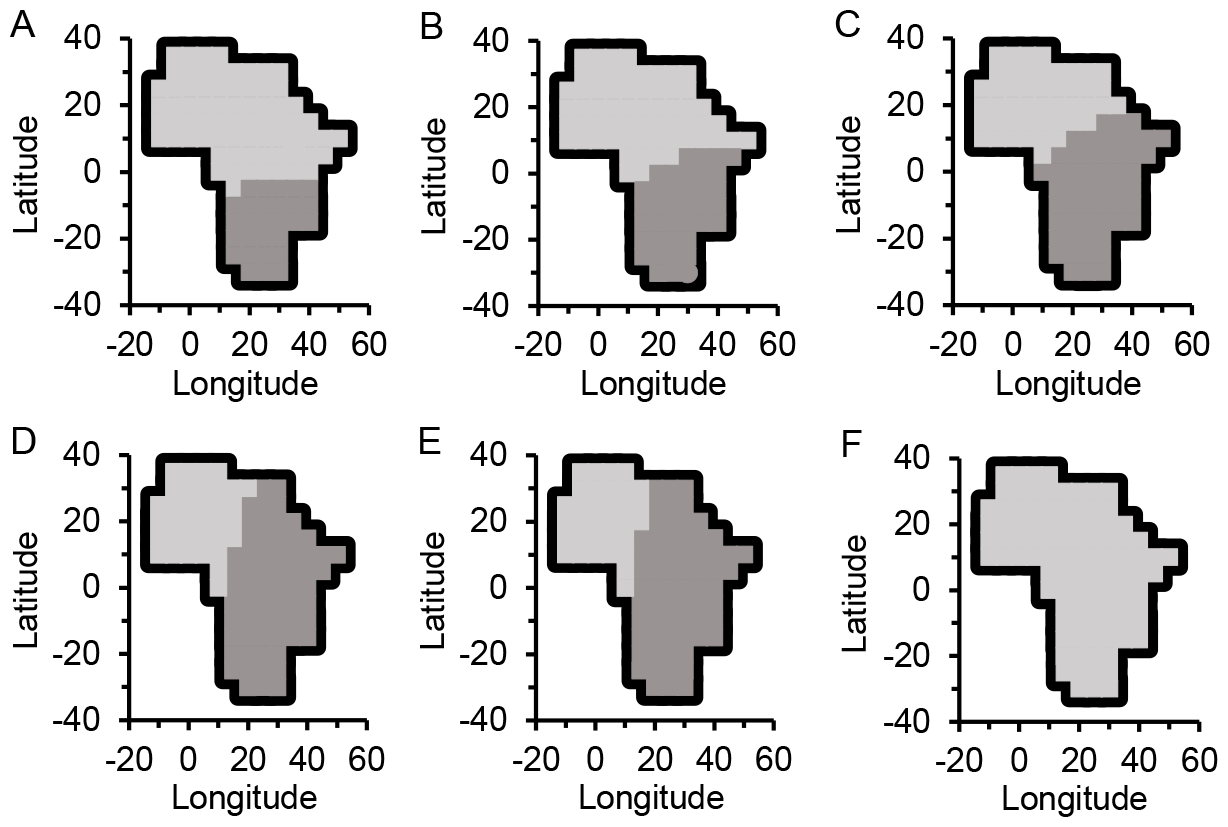
Worldwide: Y-Chromosomal Microsatellite-Derived Measures and the Origin of the Expansion. *Note*. 2A and 2D refer to TMRCA, 2B and 2E are for effective population size, whilst both 2C and 2F are with respect to expansion time. Areas in the top row (2A-2C) were found when including San in calculations, whilst the bottom row (2D-2F) was found without San. The suggested area of origin corresponds to the darker grey. With San, the area appears to be exclusively in Africa regarding TMRCA, effective population size, and expansion time. Without San, the area is partially in Africa for TMRCA and effective population size, and not at all in Africa for expansion time. With San, peak points were (30°S, 25°E) for TMRCA, and (30°S, 30°E) both for effective population size and for expansion time. Without San, the peak point for TMRCA was at Teita coordinates from von Cramon-Taubadel and Lycett (2008), the peak point was at (10°N, 50°E) for effective population size, and in Delhi (the Delhi coordinates found in von Cramon-Taubadel & Lycett, 2008) for expansion time.

**Figure 3.**
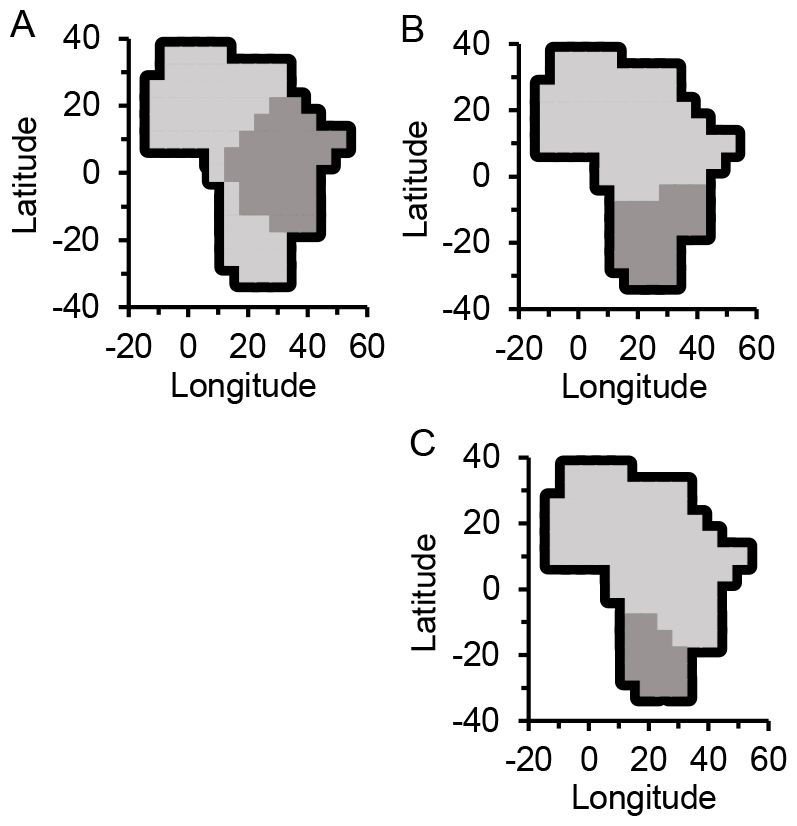
Within Africa: Y-Chromosomal Diversity and Expansion. *Note*. 3A is for repeat unit diversity, and both 3B and 3C concern haplotype diversity (3B being found including the diversity of Pygmies of Gabon presented in de Filippo et al., 2011, and 3C being without their diversity). A darker grey signifies the origin area

**Figure 4.**
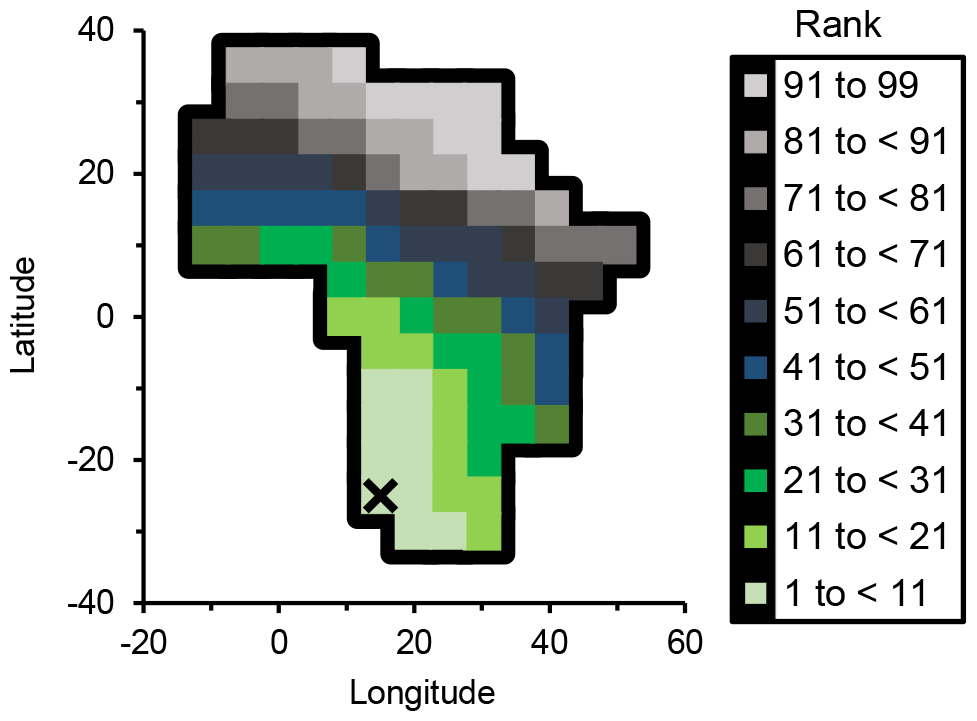
Origin of Expansion Estimated Across Genetic Diversities. *Note*. Ranks were contributed to by autosomal microsatellite heterozygosity, autosomal SNP haplotype heterozygosity, mitochondrial diversity, X-chromosomal microsatellite heterozygosity, and Y-chromosomal haplotype diversity. The location ranked at number 1 is represented with the X. The south of Africa (Choudhury et al., 2021) is where the origin appears to be.

### Geographical distance

When new geographical distances were calculated (e.g., regarding diversity in de Filippo et al., 2011), they were sought using Williams (2011). Distances were found from origin locations (Betti et al., 2013; Tishkoff et al., 2009; von Cramon-Taubadel & Lycett, 2008) to coordinates found in de Filippo et al. (2011), and between the coordinates in de Filippo et al. (2011). Naturally, 0 km was the geographical distance employed with respect to how far away from itself a population was situated. As for the Colombian and Mayan grouping of Shi et al. (2010), distances between the combined grouping and each origin were obtained by averaging distances calculated in Cenac (2022) as appropriate. When testing if diversity does associate with distance, the distance used can be from the peak point (Atkinson, 2011; Cenac, 2022), and this was done in the present study.

### Correlation

Correlation tests (regarding Y-chromosomal diversity and distance from peak points) were done in R Version 4.0.5. Holm-Bonferroni corrections (Holm, 1979) were applied to *p*-values through the Gaetano (2013) spreadsheet. Like before (Cenac, 2022), using Chen (2016) and Savin and White (1977), attention was given to positive spatial autocorrelation. Through a spreadsheet (Chen, 2016), it was examined whether positive spatial autocorrelation was evident. The spreadsheet produces a spatial Durbin-Watson (*DW*), and bounds regarding Durbin-Watson can be used on the spatial *DW* (Chen, 2016), and indeed were in the current study. The bounds utilised were the 5% ones from Savin and White (1977).

## Results and discussion

### Worldwide: Y-chromosome-derived TMRCA, effective population size, and expansion time

When San featured in analyses, it was found that TMRCA, effective population size, and expansion time (Shi et al., 2010) each provided an area of origin which is just in Africa (Figure 2A-2C). As for whether there are two origins (one from Africa, and one from elsewhere), BICs did not indicate that there was an additional origin outside of the African continent.

In Shi et al. (2010), it was noted that San (who had a sample size of four in Shi et al.) had the greatest TMRCA, as well as effective population size and expansion time. Figure 3 of Shi et al. presents distances from Africa against TMRCA (their Figure 3A), expansion time (their Figure 3B), and effective population size (their Figure 3C) – a datapoint appears to be unexpectedly great for TMRCA, expansion time, and perhaps effective population size too. It is clear that this datapoint is for San (from looking at Table S1 of Shi et al., 2010, and from references to San in Shi et al.). (Without San, the correlation coefficients in Shi et al. would likely have become more negative, numerically – see Table 1.) When using the Y-chromosome-derived variables and distances from peak points,^9^ San indeed had an unexpectedly high TMRCA (*z*-scored residual of 5.03), effective population size (*z*-scored residual of 3.39), and expansion time (*z*-scored residual of 4.11).

**Table 1.** Correlation Coefficients Regarding Y-Chromosomal Microsatellite-Derived Variables and Distance From (10°N, 40°E), With and Without San.

| Variable | San present in analysis? | Correlation coefficient |
| --- | --- | --- |
| TMRCA | Present | -.45 |
|  | Absent | -.56 |
| Effective population size | Present | -.55 |
|  | Absent | -.59 |
| Expansion time | Present | -.40 |
|  | Absent | -.43 |

*Note.* Shi et al. (2010) used distances from Addis Ababa. Coordinates for Addis Ababa were found in von Cramon-Taubadel and Lycett (2008). For Table 1, the present study used distances (Cenac, 2022) from the origin (i.e., from Betti et al., 2013 or von Cramon-Taubadel & Lycett, 2008) which was nearest to the Addis Ababa coordinates. This origin was (10°N, 40°E), and it was used in place of Addis Ababa. Correlation coefficients were calculated regarding the Y-chromosome-derived variables and distance from that origin. Also see comments regarding TMRCA in Shi et al. (2010, p. 391).

When BICs were recalculated without San, origin areas were then re-estimated (Figure 2D-2F) – variables seemed less representative of expansion; areas were no longer exclusive to Africa (TMRCA and effective population size) or were fully outside of the continent (expansion time). For the origins outside Africa, only Tel Aviv (von Cramon-Taubadel & Lycett, 2008) was inside the origin area for TMRCA and effective population size. Therefore, regarding both of these Y-chromosome-derived variables, it is likely that Africa still had most of the origin area. For both variables, the presence of an additional origin (one non-African *alongside* an African one) was not supported.

For expansion time, Asia had the area either entirely, or some of it – from the origins used, the peak point was at von Cramon-Taubadel and Lycett (2008) coordinates for Delhi like with Y-chromosomal heterozygosity in Cenac (2022) and none of the other origin locations were inside four BICs of it. Using methodology from preceding research (Manica et al., 2007), it was seen if adding distance from any of the African origins improved model fit (thereby indicating an African origin too) according to BICs (same threshold used as presumed to have been utilised in Manica et al., 2007); there was no support.^10^

Therefore, it does not seem clear if expansion from Africa is echoed in Y-chromosomal microsatellite-derived TMRCA and effective population size, and perhaps the expansion would not seem to be signaled in the expansion time (cf. Xu et al., 2015) despite TMRCA, effective population size, and expansion time being associated with distance from Africa in Shi et al. (2010). Whether San were included or not, BICs were not indicative of quadratic relationships regarding distance and either TMRCA, effective population size, or expansion time.

### Within Africa: Y-chromosomal diversity

Correlation tests were run and origin areas were estimated regarding diversity inside Africa. These analyses used repeat unit diversity as well as haplotype diversity which de Filippo et al. (2011) calculated from Y-chromosomal microsatellites. Repeat unit diversity declined linearly as distance from the peak point (0°N, 25°E) increased, *r*(24) = -.45, *p* = .043, spatial *DW* = 2.04. This diversity suggested an origin area (Figure 3A) which covered the Tishkoff et al. (2009) Africa-exit waypoint. Therefore, it is questionable if the repeat unit diversity suggests expansion from Africa. Incorporating distance from the Tishkoff et al. (2009) waypoint into the strongest model gave no improvement on the prediction of repeat unit diversity, therefore not supporting an additional origin.

On the basis of BICs, haplotype diversity was unable to suggest an origin, and so the origin was estimated merely based on the direction of correlation coefficients regarding haplotype diversity (Figure 3B). When using geographical distance from the peak point (30°S, 20°E), one of the populations (Pygmies of Gabon in de Filippo et al., 2011) had a very low haplotype diversity for their distance from the peak point (*z* = -3.35); in their absence, BICs (alongside the direction of coefficients) led to the estimation of an origin area (Figure 3C), and a negative, linearly correlation was found with distance from the peak point (25°S, 15°E), *r*(23) = -.44, *p* = .043, spatial *DW* = 2.29. The Tishkoff et al. (2009) waypoint was not in the origin area, which (given Figure 3C) indicated that the origin area for haplotype diversity was specific to Africa. Moreover, the addition of the distances from the Tishkoff et al. (2009) waypoint (to the strongest model) was of no aid in predicting haplotype diversity, and therefore haplotype diversity was not found to support an additional origin outside Africa or/and expansion into Africa. The origin areas for repeat unit diversity and haplotype diversity are generally in disagreement, overlapping only slightly (Figure 3A and 3C). Dissimilarity between origin areas has been observed previously for different diversities (e.g., autosomal diversity and cranial diversity) (Manica et al., 2007).

Autosomal microsatellite heterozygosity within Africa seems like it may deliver an insight into where the worldwide expansion started (Cenac, 2022). Autosomal microsatellite heterozygosity, across populations worldwide, lessens as distance from Africa advances (e.g., Prugnolle et al., 2005) and supports an African origin for expansion (Manica et al., 2007), and indeed, within Africa, this diversity falls whilst distance from the south increases, and indicates that expansion arose in southern Africa – autosomal microsatellite heterozygosity inside Africa may indicate the origin of expansion (Cenac, 2022).^11^ Therefore (also given Footnote 2), results in the present study raise the possibility of Y-chromosomal microsatellite haplotype diversity *within* Africa indicating the expansion regardless of whether haplotype diversity worldwide indicates an African origin or not. Hence, southern Africa (Choudhury et al., 2021) would seem to be the region of Africa which the haplotype diversity puts forward to be the origin of expansion.^12^ An estimate for where the origin started using genetic and cranial data did also point to the south (Cenac, 2022), and is thereby supported by the haplotype diversity. Nonetheless, by no means do all evaluations (of where the origin is) indicate the southern region (Ray et al., 2005).

### Overall origin estimated from genetic diversities

As Y-chromosomal haplotype diversity appeared like it was indicative of expansion, this haplotype diversity was included alongside four genetic diversities when forming a broad estimate of where the expansion started out from using an approach based on averaging ranks (as detailed in *Method*). These four genetic diversities were autosomal microsatellite heterozygosity (Pemberton et al., 2013), autosomal SNP haplotype heterozygosity, mitochondrial diversity, and X-chromosomal microsatellite heterozygosity (Balloux et al., 2009). Southern Africa (Choudhury et al., 2021) appeared as the strongest contender for being the origin (Figure 4).

## Conclusion

The Y-chromosomal microsatellite-derived variables of particular interest in the present study were TMRCA, effective population size, expansion time, haplotype diversity, and repeat unit diversity. Of them, only haplotype diversity emerged as clearly having congruency with expansion from Africa. Moreover, the heterozygosity of Y-chromosomal microsatellites does not seem to generally convey expansion from Africa (Cenac, 2022; Figure 1). And so, not only Y-chromosomal microsatellite heterozygosity (Cenac, 2022) but Y-chromosomal microsatellite-derived variables more broadly may give the impression of not generally being representative of the expansion. However, previous research (Cenac, 2022) appears to imply that Y-chromosomal microsatellite heterozygosity inside of Africa is potentially agreeable with the expansion (see Footnote 2). Also, preceding studies are not known to have seen whether TMRCA, effective population size, or expansion time suggest an Africa-only area of origin, whether with respect to the Y chromosome or otherwise. Therefore, the potential effectiveness of those three measures for indicating the area of origin is questionable. Nonetheless, through diversity, Y-chromosomal microsatellites can make a contribution to finding the African regional origin of worldwide expansion (Figure 3C) – a general estimation based on genetic diversities, inclusive of Y-chromosomal haplotype diversity, points toward the south (Figure 4).

## Footnotes

1 Attention has been given to movement other than expansion from Africa, for example, movement into the African continent (Hodgson et al., 2014) and the Bantu expansion (de Filippo et al., 2011).

2 Moreover, the Y-chromosomal microsatellite heterozygosity of populations globally (controlled for distance from Asia) presents no relationship with distance from Africa (Cenac, 2022). Therefore, Y-chromosomal microsatellite heterozygosity would not appear to be communicative of expansion from Africa (Cenac, 2022). Assuming that the expansion started in Africa (e.g., Manica et al., 2007; Ramachandran et al., 2005), a diminishing of diversity (with broadening distance from Africa) is harmonious with expansion from Africa (e.g., Betti et al., 2009; Manica et al., 2007; Ramachandran et al., 2005); with respect to the negative coefficient inside Africa regarding Y-chromosomal microsatellite heterozygosity (Cenac, 2022) – Cenac (2022) seems to be implying that the coefficient could give some indication of expansion from Africa being hinted at in Y-chromosomal microsatellite heterozygosity within Africa.

3 SNP stands for single nucleotide polymorphism (e.g., Balloux et al., 2009).

4 Research has compared population groupings in terms of their Y-chromosomal diversity (de Filippo et al., 2011).

5 De Filippo et al. (2011) did not find an indication of a decrease of Y-chromosomal diversity (referring to certain haplogroups) amongst Bantu language populations. However, they do refer to a trend in Pereira et al. (2002); whilst Pereira et al. (2002) do make reference to a geographical trend in haplotype diversity, they used Y-chromosomal haplotype diversities of just seven populations (the same number as the within-Africa Y-chromosomal heterozygosity coefficient in Cenac, 2022).

6 In Tishkoff et al. (2009), the waypoint is for expansion from Africa to Asia (Asia clearly being right next to Africa in maps in Tishkoff et al., 2009).

7 To calculate *r* correlation coefficients and *R*^2^s, distances calculated in Cenac (2022, 2023) were used apart from with respect to Y-chromosomal haplotype diversity.

8 From peak points for each of three types of autosomal diversity, Cenac (2022) calculated a centroid as an overall peak point for autosomal diversity (which was then used in the estimation of the more general centroid calculated across variables as a pointer to where the expansion started). Moreover, that research used the same autosomal SNP, X-chromosomal, and mitochondrial data as the present study when calculating the general centroid and evaluating the general geographical distribution of peak points.

9 Southern Africa (Choudhury et al., 2021) is where the peak points (Figure 2) were.

10 Regarding previous research in which Y-chromosomal microsatellite heterozygosity pointed to an origin external to Africa, that research examined whether an African origin was suggested (in the presence of an Indian origin), but using a different method (Cenac, 2022) than the present study.

11 However, unlike autosomal microsatellite heterozygosity for example (e.g., Manica et al., 2007; Prugnolle et al., 2005), it is not known if, amongst populations globally, Y-chromosomal microsatellite haplotype diversity declines with the spreading of distance from Africa (although, see Hammer et al., 2001), and points to an area of origin utterly within Africa.

12 Origin coordinates (on the inland boundary of the origin area) were checked to see which country they are in using Bartholomew illustrated reference atlas of the world (1985) – this was to see if they were within the southern African definition employed in Choudhury et al. (2021).

